# The influence of somatosensory and auditory evoked potentials on concurrent transcranial-magnetic stimulation – electroencephalography recordings

**DOI:** 10.1101/2021.11.17.469035

**Authors:** Nahian S Chowdhury, Nigel C Rogasch, Alan Chiang, Samantha K Millard, Patrick Skippen, Wei-Ju Chang, Katarzyna Bilska, E. Si, David A Seminowicz, Siobhan M Schabrun

## Abstract

**Background:** Transcranial magnetic stimulation (TMS) evoked potentials (TEPs) can be used to index cortical excitability. However, it remains unclear to what extent TEPs reflect somatosensory and auditory-evoked potentials which arise from the scalp sensation and click of the TMS coil, as opposed to transcranial stimulation of cortical circuits.

**Objectives:** The present study had two aims; a) to determine the extent to which sensory potentials contaminate TEPs using a spatially matched sham condition, and b) to determine whether sensory potentials reflect auditory or somatosensory potentials alone, or a combination of the two.

**Methods:** Twenty healthy participants received active or sham stimulation, with the latter consisting of the click of a sham coil combined with scalp electrical stimulation. Earplugs/headphones were used to suppress the TMS click noise. Two additional control conditions i) electrical stimulation alone and ii) auditory stimulation alone were included in a subset of 13 participants.

**Results:** Signals from active and sham stimulation were correlated in spatial and temporal domains, especially >70ms post-stimulation. Relative to auditory or electrical stimulation alone, combined (sham) stimulation resulted in a) larger evoked responses b) stronger correlations with active stimulation and c) a signal that could not be explained by the linear sum of electrical and auditory stimulation alone.

**Conclusions:** Sensory potentials can confound data interpretations of TEPs at timepoints >70ms post-TMS, while earlier timepoints appear reflective of cortical excitability. Furthermore, contamination of TEPs cannot be explained by auditory or somatosensory potentials alone, but instead reflects a non-linear interaction between both sources. Future studies may benefit from controlling for sensory contamination using sham conditions that are spatially matched to active TMS, and which consist of combined auditory and somatosensory stimulation.

## Introduction

Transcranial magnetic stimulation (TMS) produces evoked potentials (TEPs) during electroencephalography (EEG) that can index cortical excitability [1-3]. While traditional assessment of excitability involves TMS delivered to motor cortex with output measured from peripheral muscles, TMS-EEG permits assessment of cortical excitability not confounded by spinal/peripheral excitability, and the assessment of excitability from non-motor regions [3]. Although TMS-EEG holds promise in exploring functional brain activity and connectivity, TEPs can be contaminated by sensory input arising from the TMS pulse, confounding data interpretation.

Two kinds of sensory-evoked potentials may contribute to TEPs: auditory potentials elicited by the “clicking” of the TMS coil, and somatosensory potentials elicited by the “flicking” sensation on the skin. Several masking methods have been used to suppress these sensory inputs [4-7]. For example, white noise played through headphones has been used to mask the click sound [1, 8], while a thin layer of foam placed between the TMS coil and EEG cap has been used to minimize the scalp sensation [9]. However, recent studies have shown that even when these methods are used, sensory contamination of TEPs is still present [4-6]. Specifically, TMS produced similar signals to sham conditions that mimicked the auditory and/or somatosensory aspects of active TMS (e.g., scalp electrical stimulation with a TMS click away from the scalp). This suggests masking may not be sufficient to prevent sensory contamination, leading authors [4, 5] to recommend the use of sham conditions to control for sensory contamination.

One limitation of previous sham conditions is the sensory components were not spatially matched to active TMS. For example, four studies induced auditory potentials by delivering active TMS at a distance from the scalp, and concurrently administered mild scalp electrical stimulation to induce somatosensory potentials [4, 7, 10, 11], while another study used a sham condition where active TMS was delivered to the shoulder [5]. These studies typically demonstrated sensory contamination between ∼60-250ms post-stimulation, with earlier timepoints (<60ms) less impacted by sensory contamination [5, 6, 10, 12]. However, the use of sham conditions that do not induce auditory or somatosensory potentials that are spatially matched to active TMS may underestimate the degree of sensory contamination.

Another unaddressed question is the extent to which contamination of TEPs is explained by auditory or somatosensory potentials, or a combination of both. A study by Rocchi and colleagues [6] showed somatosensory contributions to TEPs were significantly smaller than auditory contributions, with auditory stimulation alone generating similar signals to active TMS ∼100-200ms after the TMS pulse. The authors concluded sensory contamination of TEPs is explained mostly by auditory potentials and controlling for these alone may be sufficient to index excitability. However, the authors did not assess the combined effects of auditory and somatosensory stimulation, as occurs in active TMS. While somatosensory contributions to TEPs are smaller than auditory contributions, the combination of these may result in evoked potentials that more strongly resemble the sensory potentials produced by active TMS. Indeed, studies have shown that two separate sensory stimuli can interact in a non-linear manner to produce larger evoked potentials [13, 14]. If this occurs in active TMS, sham conditions consisting of auditory input alone will not be sufficient to control for sensory contamination of the TEP.

The present study had two aims. The first was to determine whether similar contamination of TEPs would be observed to previous studies [5, 6, 10, 12] when using a spatially matched sham condition. The sham stimuli consisted of concurrent scalp electrical and auditory stimulation (sham coil click) over the left primary motor cortex, delivered at the same location as active TMS. The second aim was to determine the degree of TEP contamination explained by auditory or somatosensory potentials alone and in combination. This aim was addressed by including two control conditions (electrical and auditory stimulation alone) in a subsample of participants. The combination of auditory and electrical stimulation was hypothesised to produce a) larger evoked potentials, b) stronger contamination of TEPs and c) a signal that could not be explained by a linear sum of the responses from electrical or auditory stimulation alone.

## Methods

### Participants

20 healthy participants (13 males, 7 females, 16 TMS-naïve, age; 28.1 ± 5.3). Participants completed a TMS safety screen [15]. Participants were excluded if they were pregnant, reported a history of neurological or psychiatric conditions, or were taking psychoactive medication. Procedures adhered to the Declaration of Helsinki and were approved by the human research ethics committee of UNSW (HC200328). All participants provided informed written consent.

### Experimental Protocol

Figure 1 shows the protocol. Participants were seated comfortably in a shielded room. They viewed a fixation cross to minimise eye movements and did not observe changes in the setup between conditions. Participants wore both foam earplugs and headphones to reduce any potential discomfort from, and to dampen the noise from, the TMS click. Masking noise was not played through the headphones as the aim was for participants to perceive the TMS click to compare loudness ratings between conditions. The active and sham TMS coils were covered in a layer of foam (5mm thickness) to minimize any sensations related to coil vibration [9].

**Figure 1.**
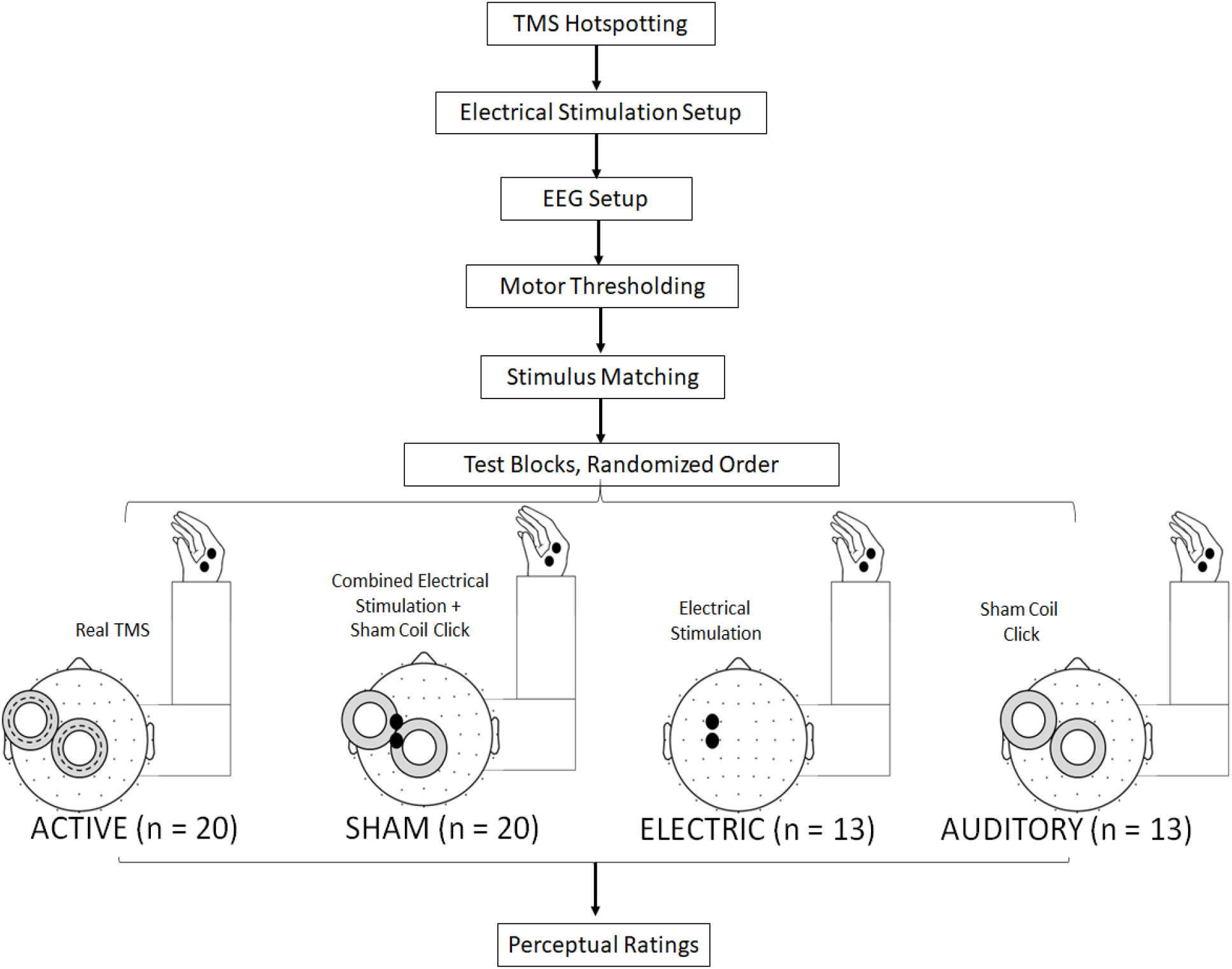
Schematic of the experimental protocol.

The experiment consisted of a single session in which participants experienced multiple blocks of TMS pulses (in randomized order), each consisting of ∼60 trials. All 20 participants received one block of 1) real stimulation over M1 (active condition) and 2) concurrent scalp electrical stimulation and a sham coil click, with both delivered over the same location as the active condition (sham condition). Inclusion of these conditions addressed the first aim of the study, which was to assess sensory contamination of TEPs using a spatially-matched sham condition. A subsample of the participants (Participants 8-20) received two additional blocks of 3) electrical stimulation applied over M1 (electric condition), and 4) auditory stimulation (sham coil click) applied over M1 (auditory condition). Inclusion of these conditions addressed the second aim of the study, which was to determine the degree of TEP contamination explained by auditory or somatosensory potentials alone and in combination.

### TMS Hotspotting

Surface electromyography (EMG) over the right first dorsal interosseous (FDI) muscle was used to record motor-evoked potentials (MEPs). EMG (8mm Ag/AgCl electrode) was recorded using NeuroMEP (Neurosoft, Russia) sampled at 2kHz. A mains filter of 50Hz was applied, with a low and high pass filter at 1000 and 10Hz respectively. Single, biphasic stimuli were delivered to left primary motor cortex using a Magstim Super Rapid^2^ Plus and a 70mm figure-of-eight air cooled coil (Magstim Ltd., UK). The coil was oriented at 45° to the midline, inducing a current in the posterior-anterior direction. The scalp site that evoked the largest MEP measured at the FDI (‘hotspot’) was determined. This location was marked and informed the location of scalp electrical stimulation.

### Electrical Stimulation

Prior to EEG setup, 8mm Ag/AgCl electrodes were placed over the scalp. In line with previous research [6], to minimize electrical stimulation artefacts in the EEG signal, the stimulating electrodes were not placed directly underneath the EEG electrodes. They were positioned in the middle of the EEG electrode cluster as close as possible to the motor hotspot. This roughly corresponded to the anode between FC1 and FC3 and the cathode between C1 and C3. To keep the electrodes in place, participants were fitted with a tight netted wig cap, which sat underneath the EEG cap. Scalp electrical stimulation was delivered using a 200 μs square wave via the Digitimer DS7AH (Digitimer Ltd., UK) with a compliance of 200V.

### Electroencephalography

EEG was recorded using a DC-coupled, TMS-compatible amplifier (ActiChamp Plus, Brain Products, Germany) at a sampling rate of 25000 Hz [7]. Signals were recorded from 63 active electrodes, embedded in an elastic cap (ActiCap, Brain Products, Germany), in line with the 10-10 system. Active electrodes result in similar TEPs (both magnitude and peaks) to more commonly used passive electrodes [16]. Recordings were referenced online to ‘FCz’ and the ground electrode placed on ‘FPz’. Electrolyte gel was used to reduce electrode impedances below ∼5kOhms.

### Motor Thresholding

TMS was delivered over the location of the stimulation electrodes. The resting motor threshold (RMT) was determined using the TMS motor thresholding assessment tool, which estimates the lowest TMS intensity required to reliably induce an MEP [17]. The test stimulus intensity was set at 110% RMT.

### Stimulus Matching

As the aim was to perceptually match the somatosensory aspects of active and sham TMS, a 2-Alternative Forced Choice task was used to determine the electrical stimulation intensity that led to a similar flicking sensation to active TMS. Participants received either electrical stimulation or active TMS in a randomized order and were asked whether the first or second stimulus led to a stronger flick sensation. The electrical stimulation intensity was then increased or decreased until participants could no longer judge the first or second stimulus as stronger. This intensity was then applied during the test blocks.

### Test Blocks

Test blocks occurred in a randomly determined order, each consisting of ∼60 pulses. 60 pulses has been shown to produce significant reliability in TEP peaks when measured within a single block [18]. All 20 participants received a single block of both active and sham stimulation. For the sham condition, an air-cooled sham coil (Magstim Ltd., UK) was simultaneously triggered with the electrical stimulation unit. The sham coil induces a small magnetic field without inducing brain current, while retaining the click sound associated with the active coil. Piloting using perceptual ratings revealed a sham stimulus intensity of 100% was required to match the loudness of the sham click as closely as possible to the active TMS click. As such, this intensity was used in the sham TMS blocks. For 13 out of the 20 participants, 2 additional blocks were included consisting of electrical or auditory (sham coil click) stimulation alone.

### Perceptual Ratings

After each block, participants were asked to rate how loud the TMS click was (0 = I did not hear anything, 10 = I heard an extremely loud click), how strong the flick sensation was (0= I felt nothing, 10 = I felt an extremely hard flick), how sharp the flick was if it was felt (0 = the flick was extremely broad, 10 = the flick was extremely narrow), how tolerable the TMS was (0 = very intolerable, 10 = very tolerable) and the extent to which they thought the brain area underneath the coil was being stimulated (0 = the brain was not stimulated at all, 10 = the brain was very much stimulated).

### Pre-processing

Pre-processing of the data was completed using EEGLAB [19] and TESA [20] in MATLAB (R2020b, The Math works, USA), and based on previously described methods [20-22]. First, bad channels were removed. The number of channels removed across participants was 2.47 ± 2.3. The period between -5 and 8.6ms (±1.83) after the TMS pulse was removed and interpolated using the ARFIT function for continuous data [23, 24]. The exact interval was based on the duration of decay artefacts. Data was epoched 1000ms before and after the TMS pulse, and baseline corrected between - 1000 and -5ms before the TMS pulse. Noisy epochs were first identified via the EEGLAB auto-trial rejection function [25] and then visually confirmed. The number of epochs excluded was 1.22 ± 2.65, 1.83 ± 2.9, 0.75 ± 1.21 and 0.83 ± 1.4 for the active, sham, electric and auditory conditions respectively. The fastICA algorithm with auto-component rejection used to remove eyeblink and muscle artefacts [10]. The mean number of components rejected was 9.4 ± 7.1, 11.3 ± 8, 12.8 ± 5 and 11.8 ± 7.5 for the active, sham, electric and auditory conditions respectively. The source-estimation noise-discarding (SOUND) algorithm was applied [21, 22], which estimates and supresses noise at each channel based on the most likely cortical current distribution given the recording of other channels. This signal was then re-referenced (to average). A band-pass (1-100Hz) and band-stop (48-52Hz) Butterworth filter was then applied. Any lost channels were interpolated.

### Statistical Analysis

#### Perceptual Ratings

JASP software (Version 0.12.2.0, JASP Team, 2020) was used to conduct Bayes paired samples t-tests (Cauchy scale = .707), comparing the active and sham conditions on ratings of perceived loudness, flick strength, flick sharpness, tolerability, and stimulation extent. To determine whether combined electrical and auditory stimulation enhanced the experience of “real” brain stimulation, stimulation extent in the sham condition was compared against the auditory and electric conditions. Bayes factors were expressed as *BF*_10_ values, where a value <=0.33 indicated evidence that the perceptual ratings were matched between conditions [26].

#### Aim 1: Sensory Contributions to TEPs

The grand-averaged signals and global mean field waveform (GMFW) were obtained for active and sham stimulation. Maxima in the GMFW were identified using the TESA peak function [20]. In line with previous studies [4, 6], based on the peaks of the GMFW of the active condition, signals were separated into early, mid and late time-points of interest (TOIs) [4, 6]. The Fieldtrip toolbox [8] was used to conduct a cluster-based permutation analysis to compare amplitude levels between active and sham conditions at each TOI. The spearman-ranked spatial correlations (across electrodes at each time-point) and temporal correlations (across time-points at each electrode) between active and sham was determined. Correlation coefficients were transformed using Fisher’s z method. For spatial correlations, the 95% confidence interval of the mean correlation value (across participants) at each time-point was assessed against zero [5]. Temporal correlations were performed separately for early, mid and late TOIs. The mean correlation value (across participants) at each electrode was assessed against zero [5]. Lastly, to determine whether the GMFW maxima in the active condition were retained after controlling for sensory-evoked potentials, signal-space projection with source-informed reconstruction (SSP-SIR) [9] was used to suppress sensory potentials observed in the sham condition from the active condition. SSP-SIR is a filter which identifies spatial commonalities between two signals and suppresses this from the target signal.

#### Aim 2: Individual Sensory Contributions to TEPs

The grand-averaged signal and GMFW were obtained for active, sham, electrical and auditory stimulation, and the spatial and temporal correlations between conditions were computed. To determine whether the combination of auditory and electrical stimulation generated stronger sensory-evoked potentials compared to each condition alone, a cluster-based permutation analysis was conducted comparing the sham condition with the electric and auditory conditions. To determine whether there were stronger correlations with active TMS when electrical and auditory stimulation was combined (sham) vs. delivered alone (electric or auditory), the spatial correlations for active-sham were compared with active-electric and active-auditory. This was done by running a two-sample t-test comparing the mean z-transformed correlation coefficients (across participants) at each time-point. Lastly, to assess whether a simple linear summation of somatosensory and auditory-evoked potentials could capture the responses observed in the sham, the responses from the electric and auditory condition were added, and spatial and temporal correlations between the summed signal and the sham signal were determined.

## Results

Two participants were excluded due to evidence of bridging across electrodes but were included in the analysis of perceptual ratings. This left 18 participants relevant to the first aim of the study, and 12 participants relevant to the second aim. Note these sample sizes remained comparable with previous studies investigating sham conditions, which ranged from 12-20 [4-6, 10]. The mean RMT was 84.0 ± 9.7 % of maximum stimulator output. The mean test electrical stimulation intensity was 5.5 ± 8.1 mA.

### Perceptual ratings

Figure 2 shows the perceptual ratings for each condition. There was no significant difference between active and sham conditions in click loudness, (*p =* 0.07, *BF*_10_ *=* 0.97), flick strength, (*p =* 0.35, *BF*_10_ *=* 0.33), flick sharpness, (*p =* 0.6, *BF*_10_ *=* 0.26) and tolerability, (*p =* 0.34, *BF*_10_ *=* 0.34). As the Bayes factors yielded inconclusive support for the matching of loudness between active and sham, a Bayesian t-test comparing loudness ratings in active and auditory conditions was conducted (n = 13). This was justified given an identical sham coil click was used in the auditory and sham conditions. This analysis yielded a BF_10_ value of .302, suggesting the click of the sham coil and active TMS were matched in terms of loudness. Participants rated active TMS as stimulating the brain to a larger extent than sham (p *=* 0.01, *BF*_10_ *=* 3.7). Stimulation extent was rated higher in the sham compared to either auditory (*p <* .001, *BF*_10_ *=* 69.5) or electrical stimulation alone (*p <* .001, *BF*_10_ *=* 782.3), suggesting the combination of auditory and electrical stimulation enhances the experience of “real” stimulation.

**Figure 2.**
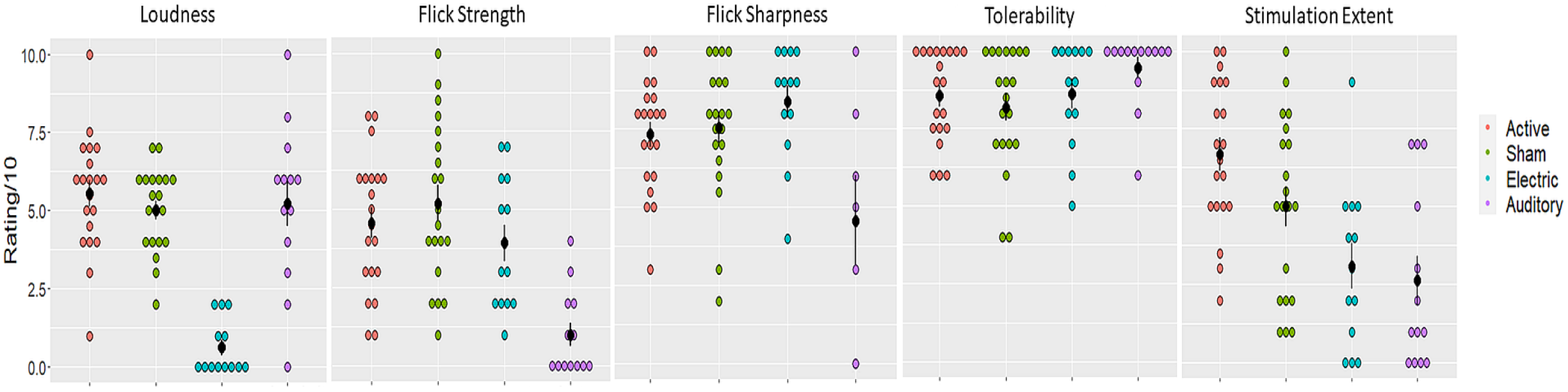
Dotplots showing participant ratings of TMS click loudness, TMS flick strength, TMS flick sharpness (which were given only if a flick was felt in the first place), tolerability and stimulation extent.

### Sensory Contributions to TMS-evoked potentials

Six maxima were identified in the active condition, including 16, 31, 43, 64, 104 and 184ms after TMS (Figure 3). Based on the peaks of the GMFW, three TOIs were selected for comparisons between conditions– early (12-70ms), middle (71-138ms) and late (139-250ms) time periods. The cluster-based permutation analysis revealed a significant negative (*p* =0.03) and positive (*p*=0.03) cluster at the early TOI (Figure 3). However, there were no significant voltage differences at mid and late TOIs (*p*’s > .05). For the suppressed signal (after SSP-SIR), the GMFW appeared substantially attenuated relative to active stimulation (Figure 3). However, besides the maxima at 43ms, all other maxima were retained. Grand-averages and scalp topographies for the active, sham, and suppressed signals for the full sample of 18, are shown in Figure 4.

**Figure 3.**
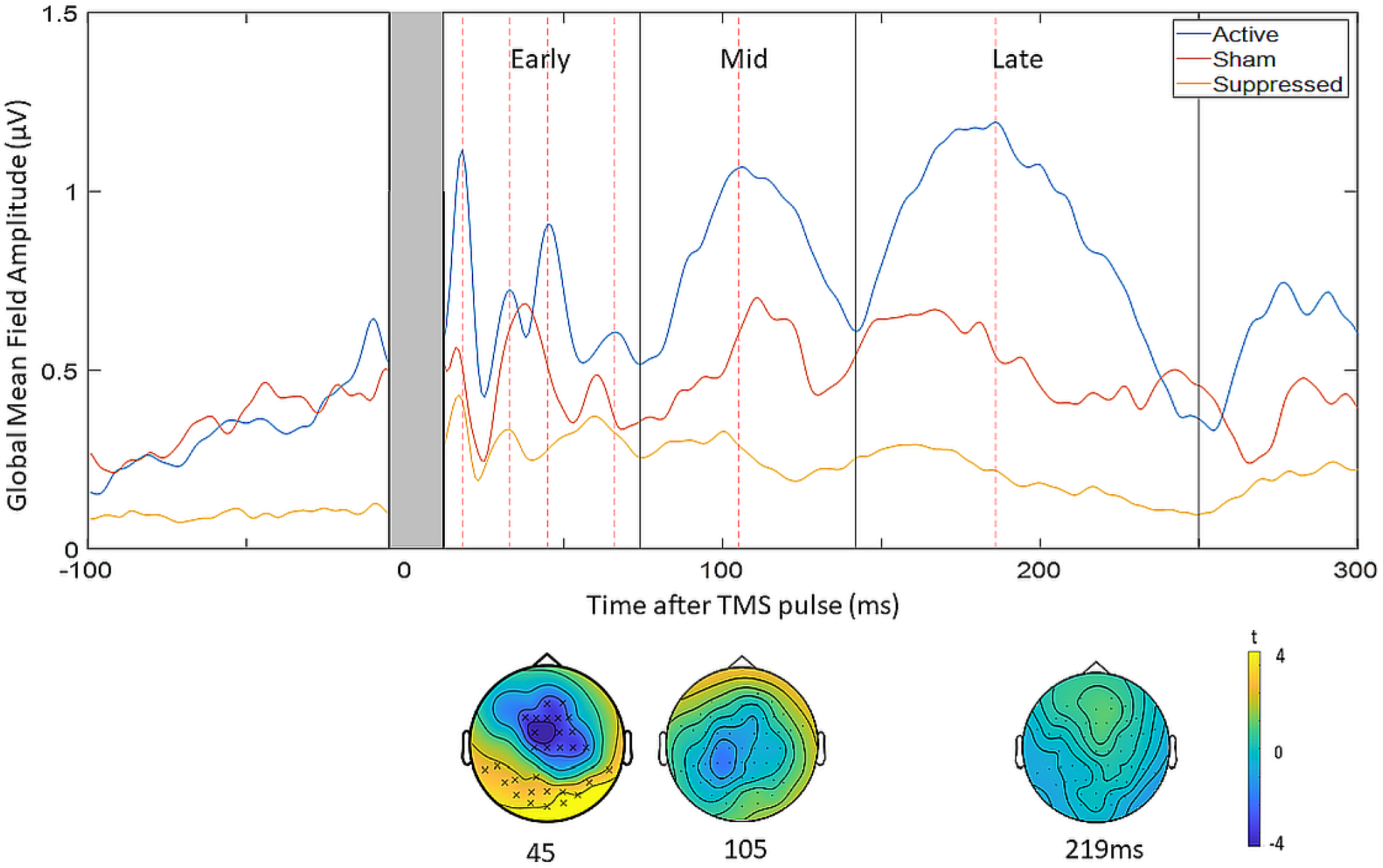
Amplitude comparisons between the active, sham and suppressed signals. The top panel shows the global mean field waveforms for the three conditions. The black solid lines represent the boundaries of the time points of interest (TOIs) – early (12-70ms), mid (71-138ms) and late (139-250ms), and red-dotted lines shows the maxima of the active signal. The bottom panel shows the cluster plots comparing the amplitude between the active and sham condition at a representative time-point at each TOI (45, 105 and 218ms). The black stars demonstrate the presence of significant positive (yellow) or negative (blue) clusters.

**Figure 4.**
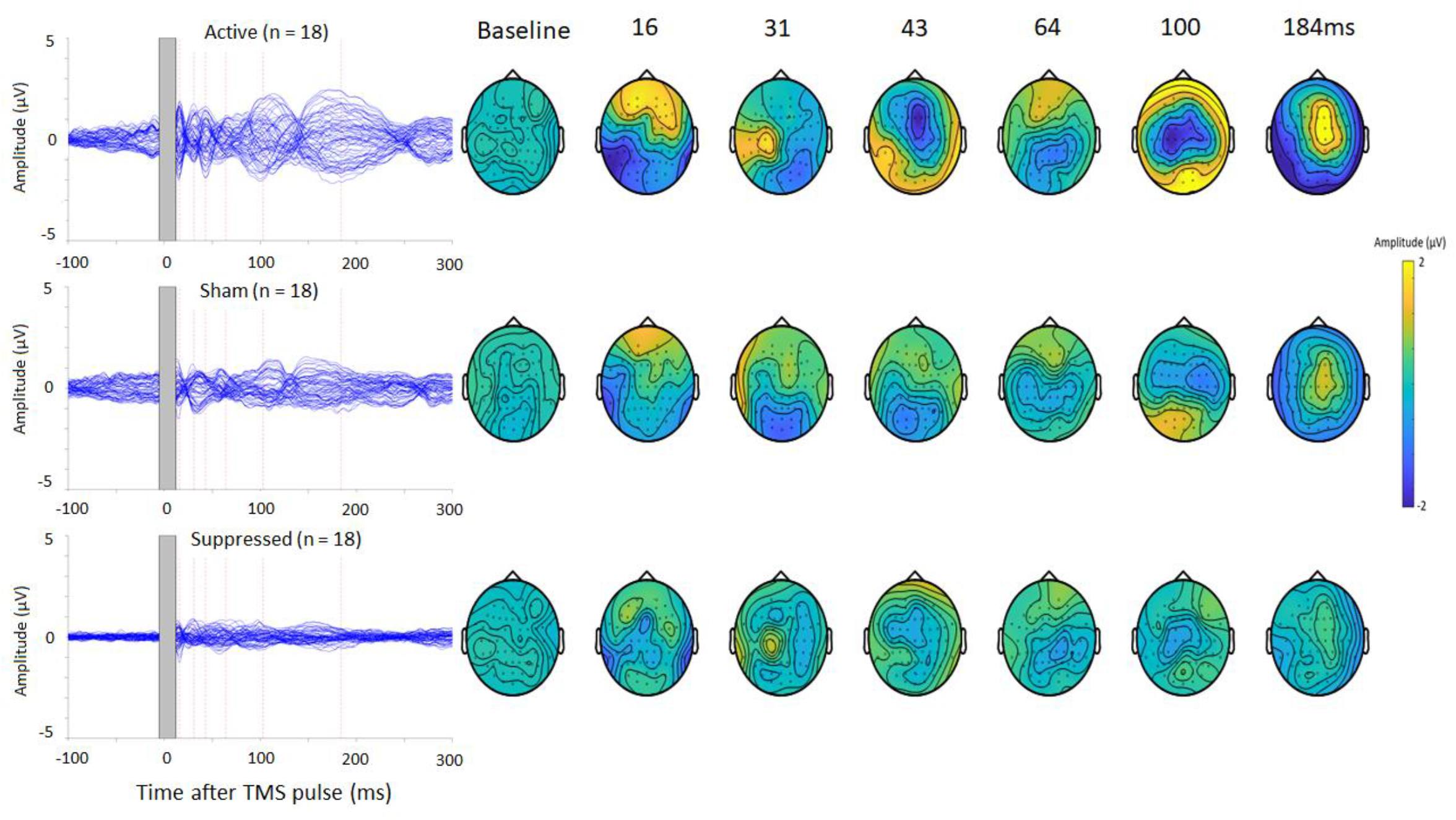
TEPs and scalp topographies following active and sham stimulation over left M1 for the full sample (n = 18). The grey shaded area represents the window of interpolation around the TMS pulse. The red dotted lines represent the time-points of the maxima from the global mean field waveforms of the active condition. The scalp topographies show the distribution of voltage at each of these time-points, and the mean topography during the baseline period.

Significant spatial correlations between the active and sham conditions were present at mid and late TOIs, but not the early TOI (Figure 5). Significant temporal correlations were present at the early (right-occipital), middle (central) and late (global) TOIs (Figure 5).

**Figure 5.**
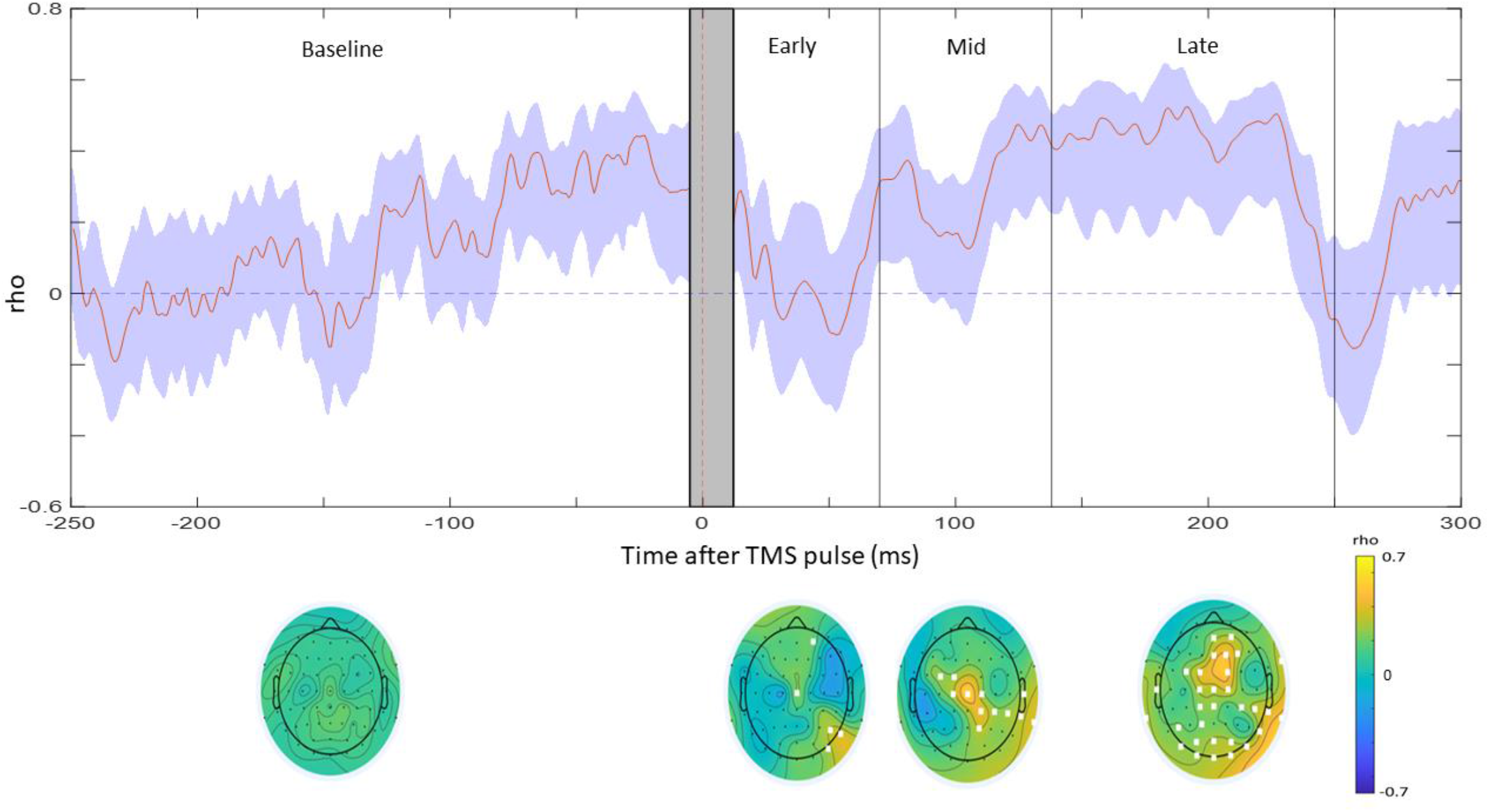
Spatial and Temporal correlations between the active and sham conditions. The top panel shows the correlations across electrodes (spatial) between 250ms before and 300ms after the TMS pulse. The blue shaded area represents 95% confidence intervals. The grey shaded area represents the window of interpolation around the TMS pulse. The black lines indicate the boundaries for the early, mid and late TOIs. The bottom panel shows heat maps of the correlations across time at each electrode, separated by time of interest (early, middle or late time-points) and during the baseline period. The white squares indicate the electrodes with significant correlations (p<.05).

### Individual Sensory Contributions to TMS-evoked potentials

Grand-averages and scalp topographies for the active, sham, electric and auditory conditions for the subsample of 12 are shown in Figure 6.

**Figure 6.**
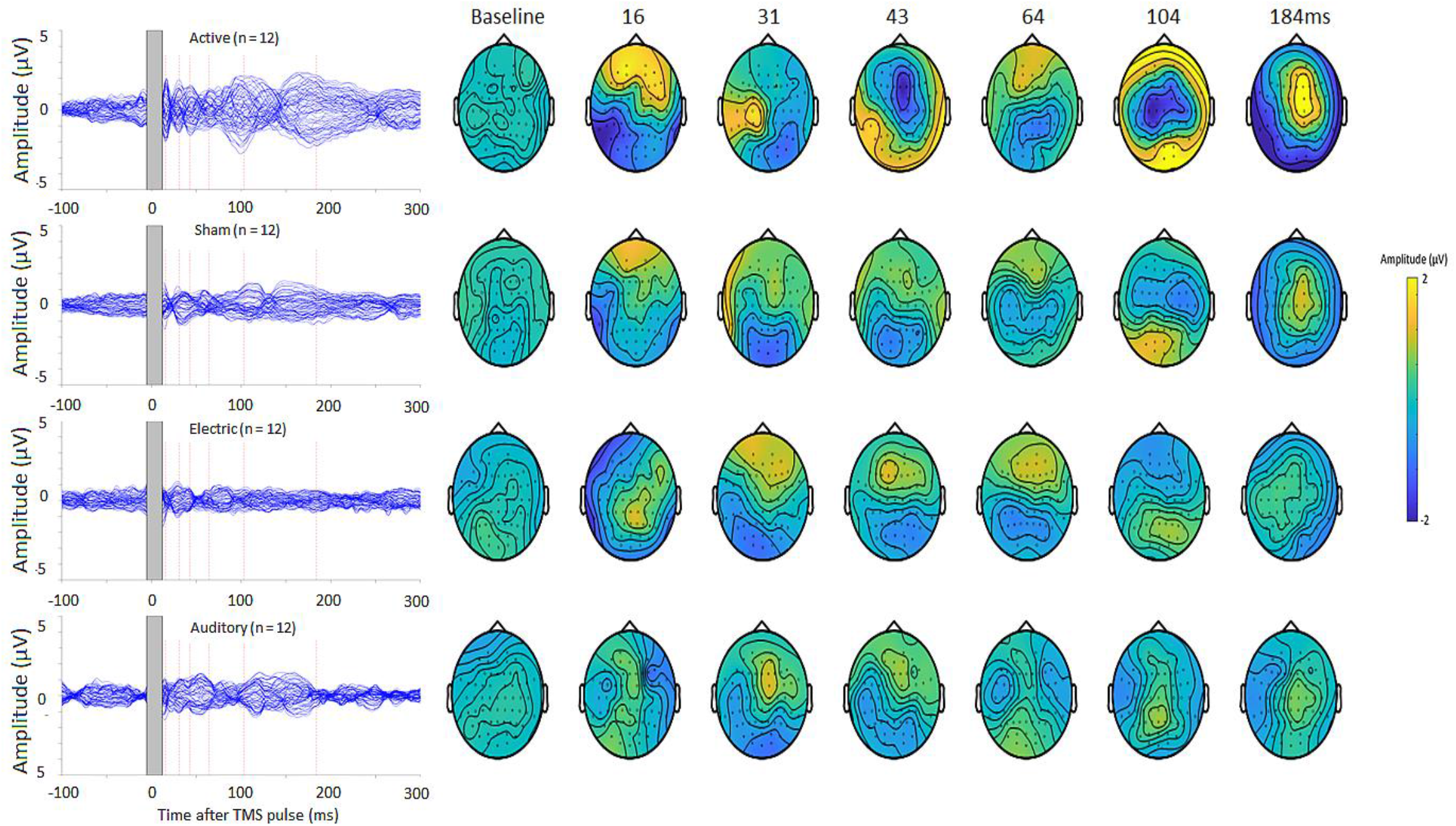
TEPs and scalp topographies following active, sham, electric and auditory stimulation over left M1 (n = 12). The grey shaded area represents the window of interpolation around the TMS pulse. The grey shaded area represents the window of interpolation around the TMS pulse. The red dotted lines represent the time-points of the maxima from the global mean field waveforms of the active condition. The scalp topographies show the distribution of voltage at each of these time-points, and the mean topography during the baseline period

The active and sham correlations (Figure 7) largely corresponded to the full sample, with significant spatial correlations at TOIs >70ms, and similarly distributed temporal correlations at all TOIs (Figure 7A). There were significant late temporal correlations between the active and electric conditions in a small subset of electrodes, but no significant positive spatial correlations (Figure 7B). When comparing the electric to the sham condition, there were significant temporal and spatial correlations at all TOIs, though these were more frequently observed at early and mid TOIs (Figure 7B). There were significant positive temporal correlations between active and auditory conditions at early (right-parietal), middle (central-occipital) and late (frontal and central-parietal) TOIs, and significant spatial correlations at mid and late TOIs (Figure 7C). There were significant temporal and spatial correlations between sham and auditory at all TOIs, though these were more frequently observed at mid and late TOIs (Figure 7C).

**Figure 7.**
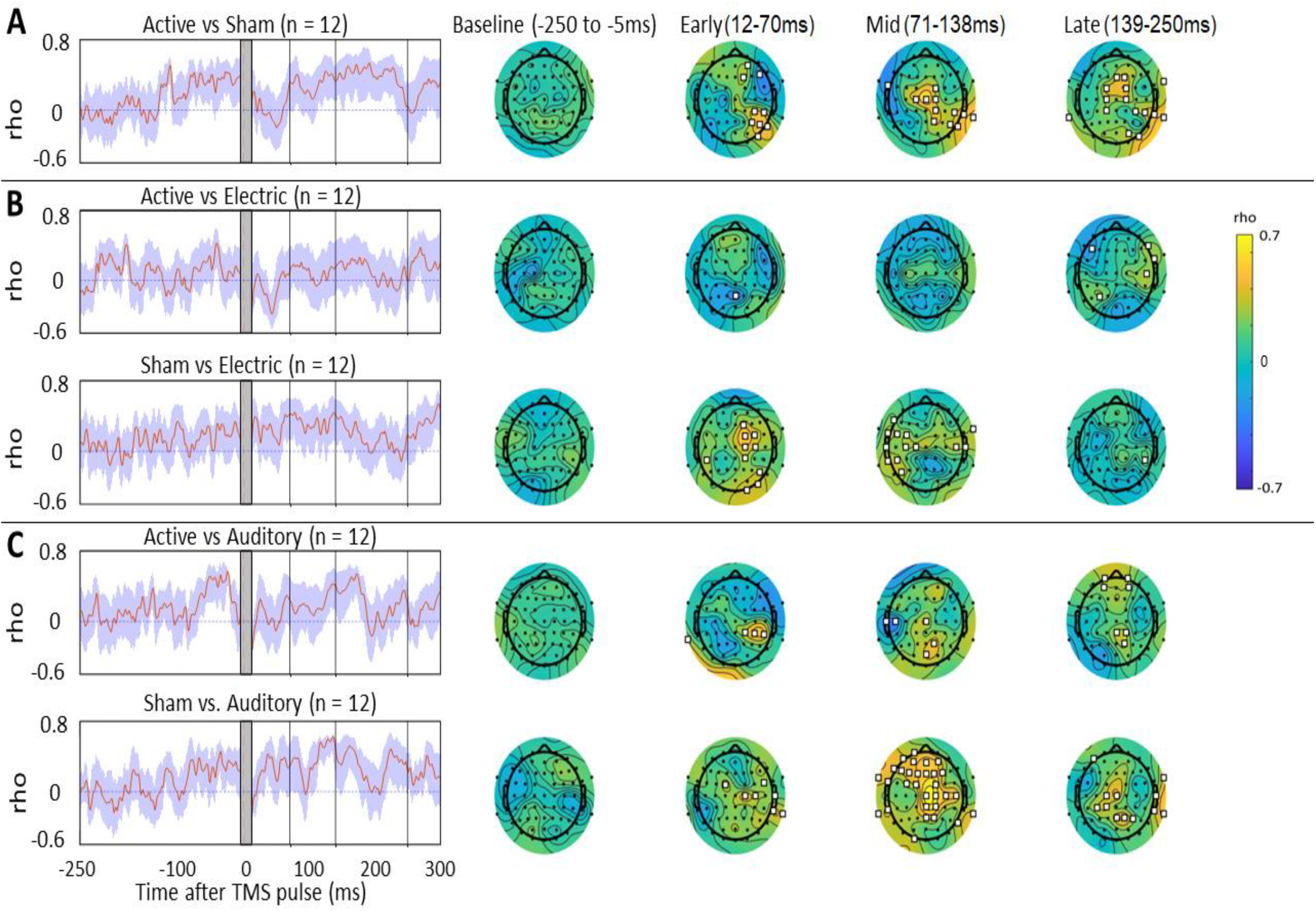
Spatial and Temporal correlations for active vs sham (panel A), active vs electric and sham vs electric (panel B) and active vs auditory and sham vs auditory (panel C). The left side shows the correlations across electrodes (spatial) between 250ms before and 300ms after the TMS pulse. The blue shaded area represents 95% confidence intervals. The vertical black lines indicate the boundaries of the early, late and mid TOIs. The grey shaded area represents the window of interpolation around the TMS pulse. The right side shows the topography maps showing correlations across time at each electrode, separated by time of interest (early, middle or late time-points) and during the baseline period. The white squares indicate the electrodes with significant correlations (*p*<.05).

Interestingly, spatial correlations in the baseline period for active-sham, active-auditory and sham-auditory were significant (Figures 5 and 7). This has been observed in previous studies [5, 6] and has been attributed to a pre-processing artefact around the TMS pulse artefact [5]. To test this possibility, the same correlation tests were conducted without the ICA pre-processing step (Figure S1). This showed spatial correlations were either weaker or non-significant in the baseline period without the ICA pre-processing step, while the correlations thereafter were preserved.

The sham-active correlations were stronger than the electric-active correlations between ∼180 and 220ms, and stronger than the auditory-active correlations between ∼180-240ms (Figure 8). To determine whether these correlation comparisons were influenced by pre-processing, the same tests were carried out without ICA (Figure S2). This showed the active-sham correlation was stronger than the electric-active correlation at the late TOI, and stronger than the auditory-active correlation at the mid TOI.

**Figure 8.**
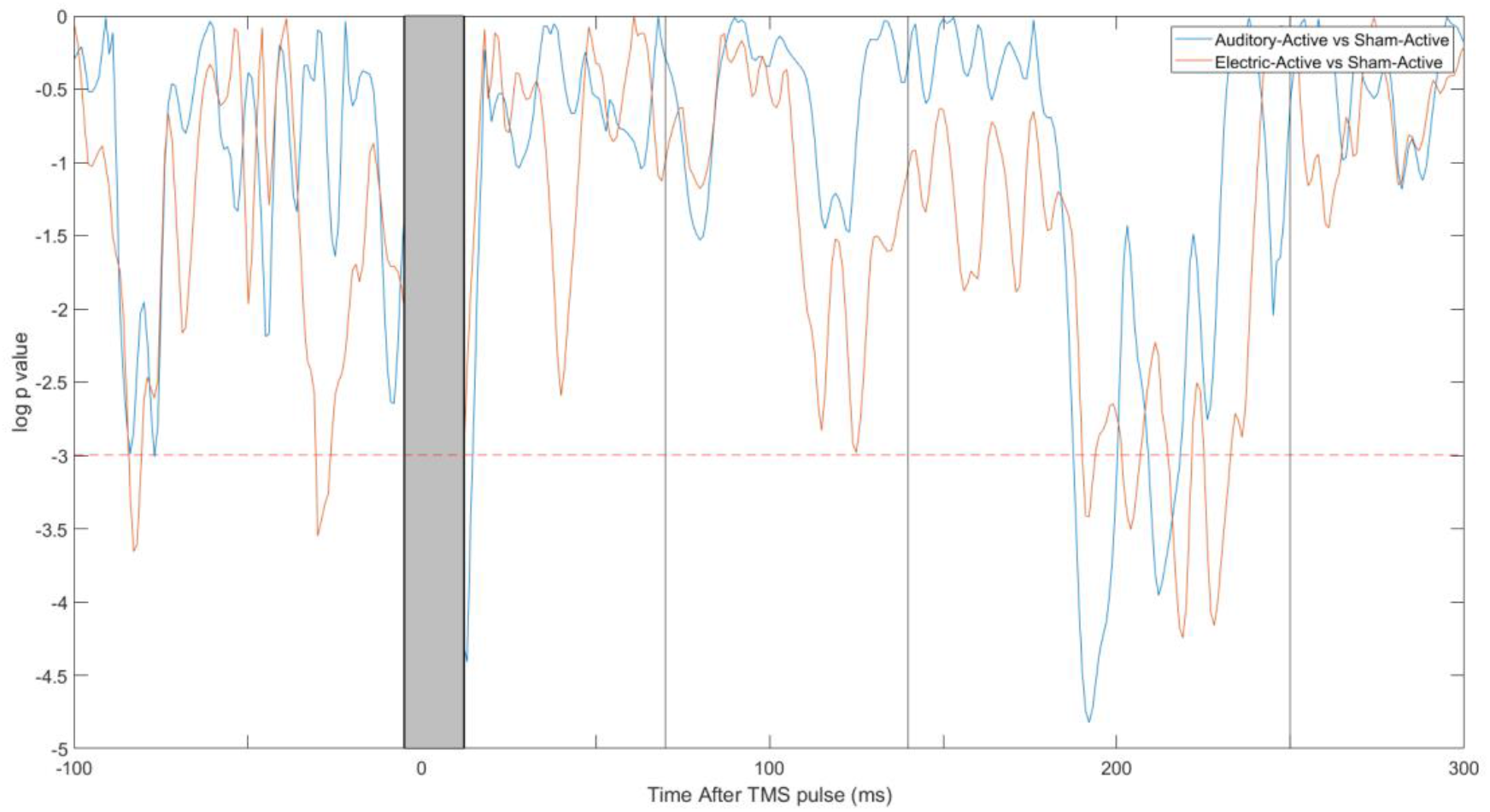
Line plots showing the statistical significance of the difference between the sham-active correlation with the auditory-active correlations (blue) and the electric-active correlations (orange). The grey shaded area represents the window of interpolation around the TMS pulse. The horizontal red dotted line shows the criteria for significance (*p =* .05). p-values were log-transformed for illustrative purposes.

The sham condition resulted in a larger evoked response compared to the electric condition in the mid and late time-points, and a larger response compared to the auditory condition in the late time-point (Figure 9).

**Figure 9.**
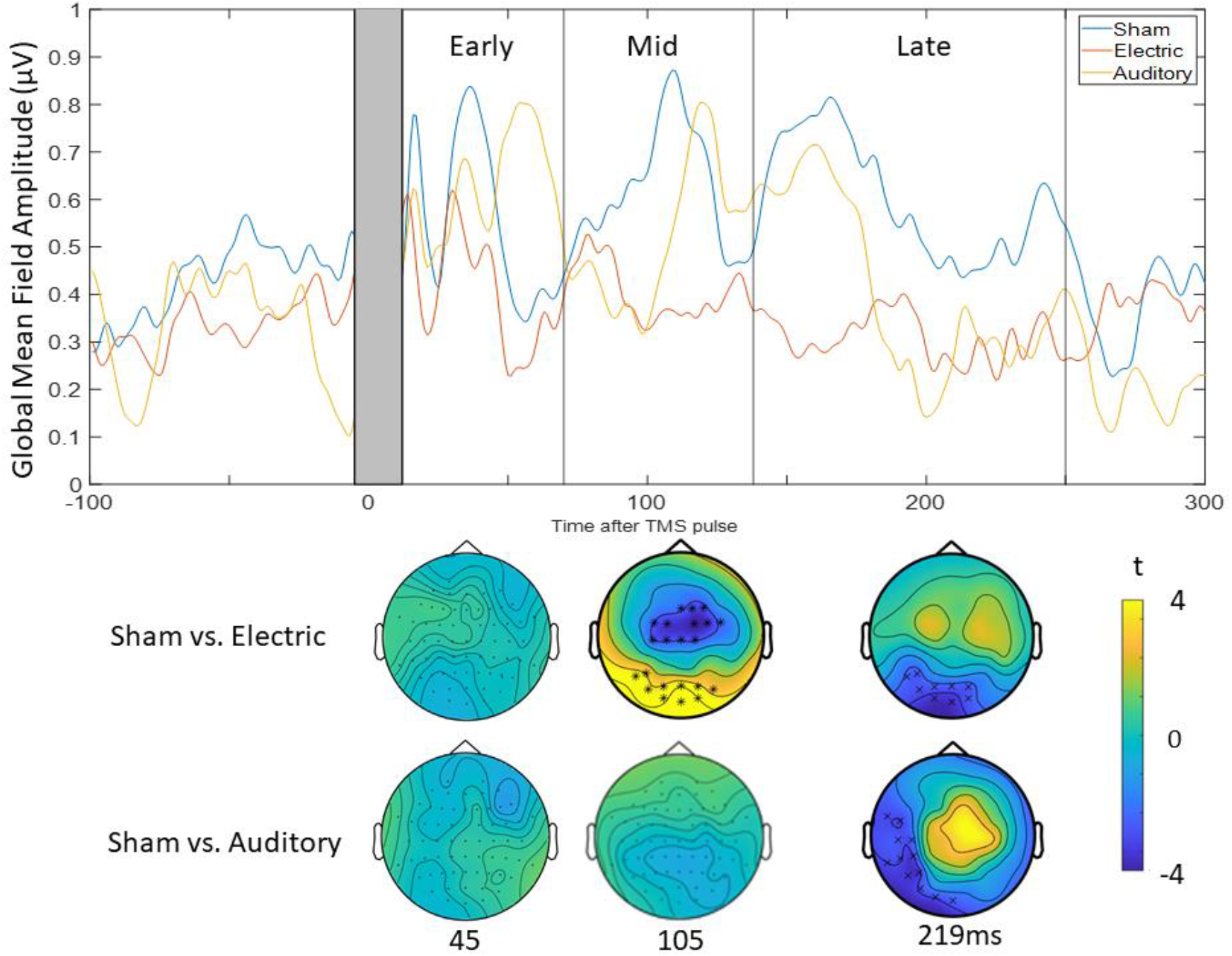
Amplitude comparisons between the sham, electric and auditory signals. The top panel shows the global mean field waveforms for the three conditions. The black solid lines represent the boundaries of the TOIs – early (12-70ms), mid (71-138ms) and late (139-250ms). The bottom panel shows the cluster plots comparing the amplitude between the sham and electric, and sham and auditory conditions, at a representative time-point at each TOI (45, 105 and 218ms). The black stars demonstrate the presence of significant positive or negative clusters.

The summed signal from the electric and auditory conditions showed high spatial and temporal correlations with the sham condition up until ∼170ms but showed relatively lower spatial correlations between 180-240ms and relatively lower temporal correlations in the late TOI (Figure 10). These findings suggest concurrent somatosensory and auditory stimuli results in a response which is not a simple linear summation of these two inputs, especially at later time-points.

**Figure 10.**
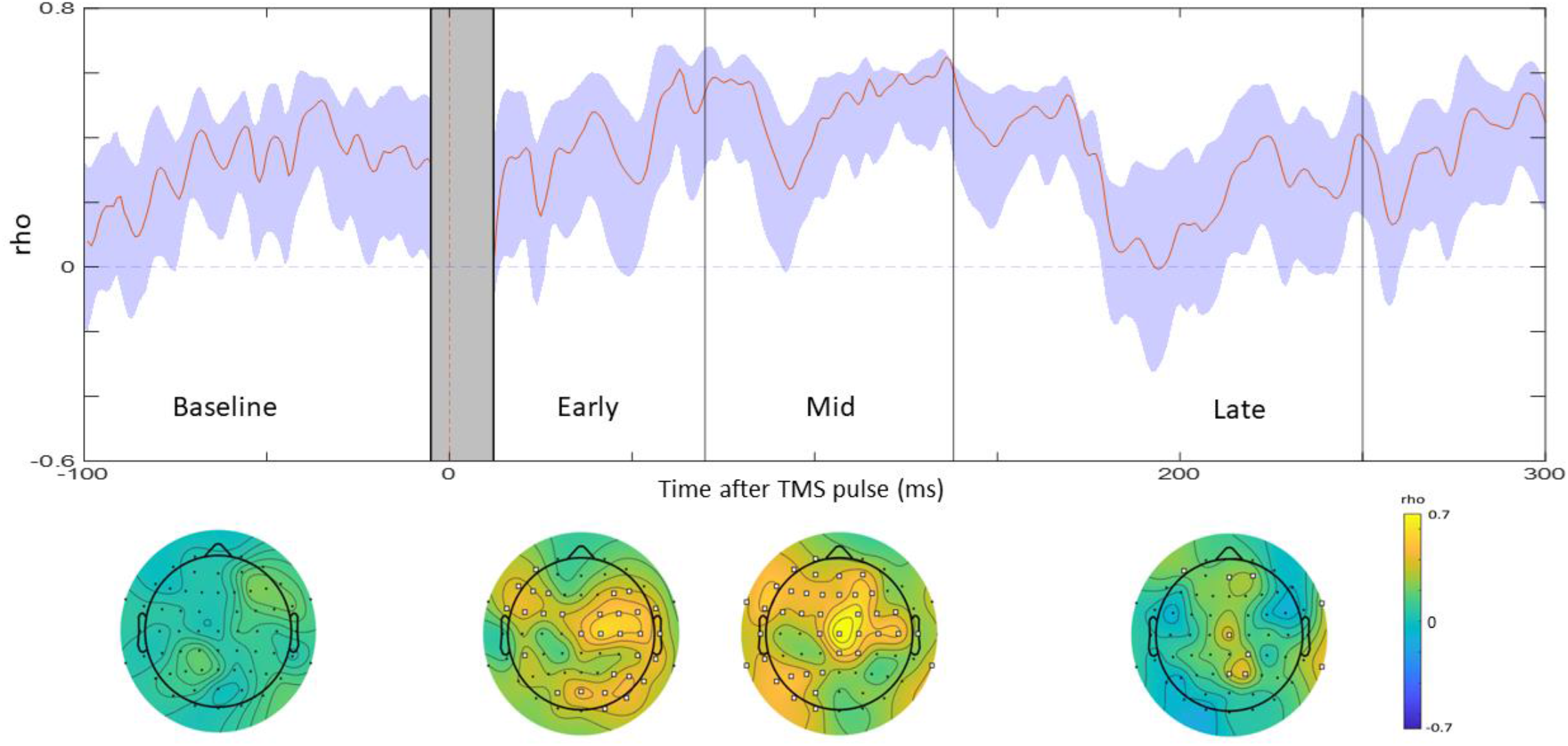
Spatial and temporal correlations between the sham condition and the summed potential of the auditory and electric condition. The blue shaded area represents 95% confidence intervals. The grey shaded area represents the window of interpolation around the TMS pulse. The left shows the correlations across electrodes (Spatial) between 250ms before and 300ms after the TMS pulse. The vertical blue lines indicate the boundaries of the early, late and mid TOIs. The right shows the maps showing correlations across time (temporal) at each electrode, separated by time of interest (early, middle or late time-points) and during the baseline period. The white squares indicate the electrodes with significant correlations (*p* <.05).

## Discussion

The first aim of the study was to determine the extent to which auditory and somatosensory-evoked potentials contaminate TEPs when using a spatially matched sham condition. Active TMS generated larger responses than sham between 12-70ms post-TMS. There were positive spatial and temporal correlations between active and sham signals at timepoints >70ms. The second aim of the study was to determine how much this contamination was explained by somatosensory or auditory potentials, or a combination of both. Relative to electrical and auditory stimulation alone, the combination of auditory and electrical stimulation (i.e., sham) resulted in a) larger evoked potentials, b) stronger correlations with active TMS and c) a signal that could not be explained by a linear sum of electrical and auditory stimulation.

### Sensory Contributions to TMS-evoked potentials

A major focus of current TMS-EEG research has been understanding how sensory-evoked potentials contaminate TEPs. Studies using sham conditions with both sub- and suprathreshold motor cortex stimulation have demonstrated that earlier timepoints (<∼60ms post-TMS) are less impacted by sensory contamination [5, 6, 10, 12]. However, the sensory aspects of most shams were not spatially matched to active TMS. In the current study, the use of a spatially-matched sham did not alter previous findings. Specifically, the amplitude of sensory-evoked potentials was significantly smaller than the active signal at earlier timepoints <70ms post-TMS. Moreover, when comparing active and sham at these earlier time-points, there were no spatial correlations, and temporal correlations existed at right occipital sites only. This replicates previous findings and support the notion that early timepoints ∼<60-70ms post-TMS represent the cortical response to TMS. While it is still unknown precisely what mechanisms the early TEP peaks represent, pharmacological studies [27] and studies comparing TEPs with single and paired-pulse MEPs [28-30] suggest early TEPs may reflect the excitability of the cortex, especially close to the site of stimulation.

An important finding was evidence of perceptual matching between active and sham conditions, particularly for the somatosensory ratings (flick sharpness and strength). Matching the perceptual aspects of active and sham stimulation has been a major challenge in previous studies [4, 5, 11]. However, most studies used sham conditions where active TMS was delivered at a distance from the scalp [4, 5, 10]. A recent study [11] achieved perceptual matching by delivering high intensity electrical stimulation in both active and sham conditions to render both indistinguishable. Though this was effective in blinding participants, the inclusion of electrical stimulation in the active condition may have altered the genuine TEP response. This highlights one advantage of the current setup as somatosensory ratings were matched without the need for electrical stimulation during active TMS. However, a more thorough assessment is required to determine whether the sham stimuli in the present study is indistinguishable from active TMS.

### Individual Sensory Contributions to TMS-evoked potentials

The findings align with previous work showing auditory potentials make a greater contribution to TEPs when compared with somatosensory potentials [6]. However, the present study also showed the amplitude of the combined response to electrical and auditory stimulation (i.e., the sham) was larger than for either electrical or auditory stimulation alone (specifically at mid/late TOIs), resulting in stronger correlations with the active condition at late TOIs. Combined stimulation was also perceptually rated as providing stronger brain stimulation. Furthermore, summating the responses from the auditory and electrical conditions did not recover the combined EEG response at late TOIs, suggesting the concurrent input of the two sensory stimuli results in a non-linear interaction within the brain. Some authors have used sham conditions consisting of only auditory stimuli [12], and others have concluded that controlling for auditory potentials alone may be sufficient to index excitability [6]. The present findings suggest even though somatosensory contributions to TEPs are smaller than auditory contributions, sensory contamination of TEPs is best explained as a non-linear interaction between somatosensory and auditory inputs. Together, these findings suggest sham conditions using concurrent auditory and somatosensory stimuli can more accurately capture sensory contamination within TEPs, and more closely match the perceptual experience of active TMS.

## Limitations

There are several limitations to discuss. First, the electrical stimulation in the present study did not induce the facial/cranial nerve stimulation present during active TMS [31, 32]. Second, the more prominent contribution of auditory stimulation to active TMS may be explained by the absence of auditory noise masking. Indeed, a recent study [6] showed mixing white noise with specific time-varying frequencies of the TMS click through headphones results in a significantly smaller auditory response. Finally, reafferent activity from the motor-evoked response in peripheral muscles can also alter TEPs [33-35] and this was not controlled for in the current study. Further studies are required comparing the signals produced by active and sham stimulation in the presence of auditory masking, as well as using somatosensory stimulation which activates cranial nerves.

## Conclusions

The present findings replicate previous studies showing sensory potentials can significantly contaminate EEG signals at timepoints ∼>70ms post-stimulation. Further, the findings provide evidence that concurrent auditory and somatosensory input can capture sensory contamination more accurately than auditory or somatosensory input alone. Future TMS-EEG studies may benefit from controlling for sensory contamination using sham conditions that are spatially matched to active TMS and consist of combined auditory and somatosensory stimulation.

## Author Contributions

N.C. was involved in conceptualization, data curation, formal analysis, investigation, methodology, project administration, validation, visualisation, writing – original draft, writing – review and editing. N.R. was involved in supervision, software, conceptualization, formal analysis and writing – review and editing. A.C. was involved in conceptualization, formal analysis and writing – review and editing S.M. was involved in conceptualization, formal analysis and writing – review and editing. P.S.,. K.B., and E.S. were involved in writing – review and editing. D.S. was involved in resources, supervision and writing – review and editing. S.M. was involved in resources, supervision and writing – review and editing.

## Supplementary Material

**Figure S1.**
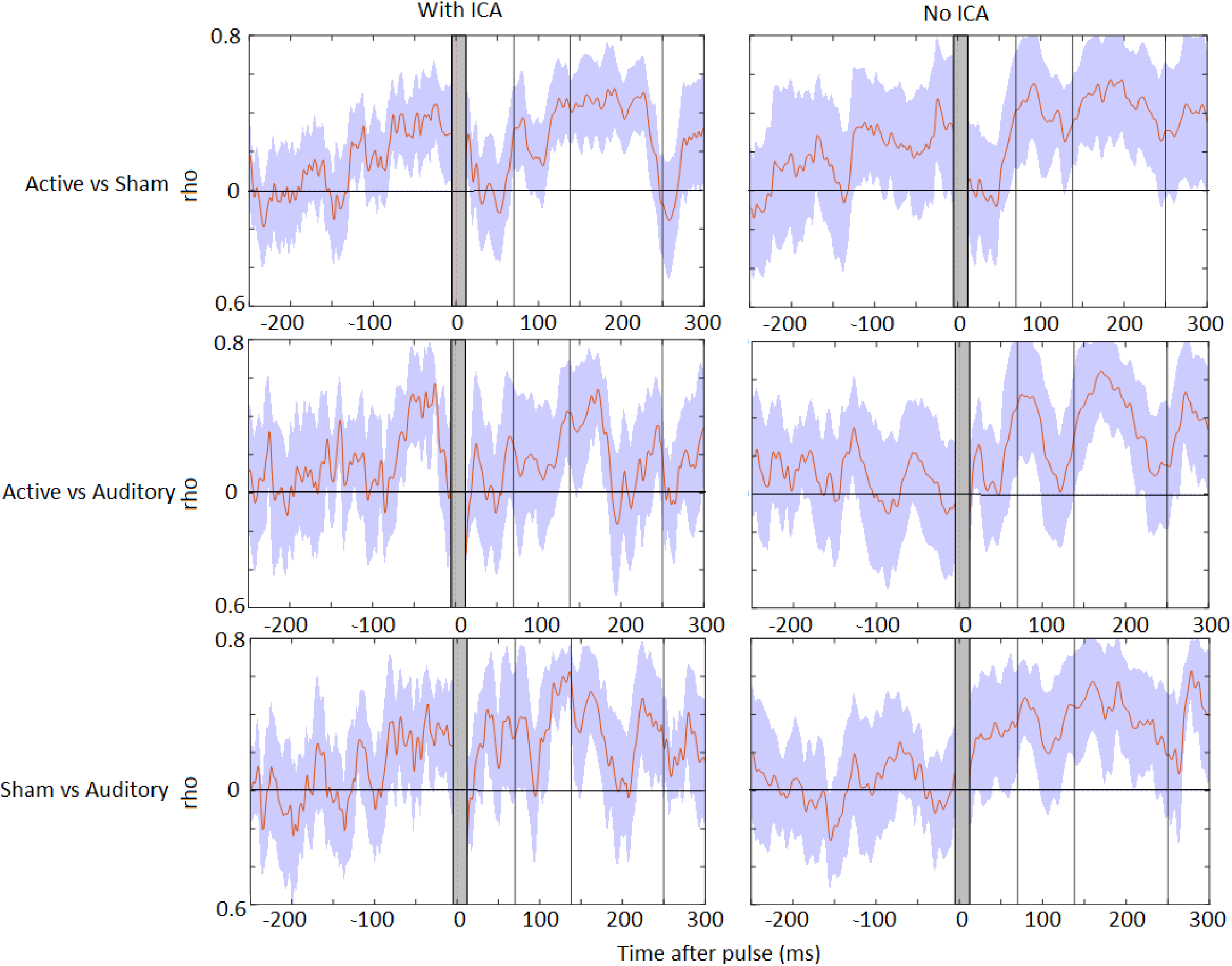
Effects of excluding ICA from the pre-processing pipeline on the Active-Sham, Active-Auditory and Sham-Auditory spatial correlations

**Figure S2.**
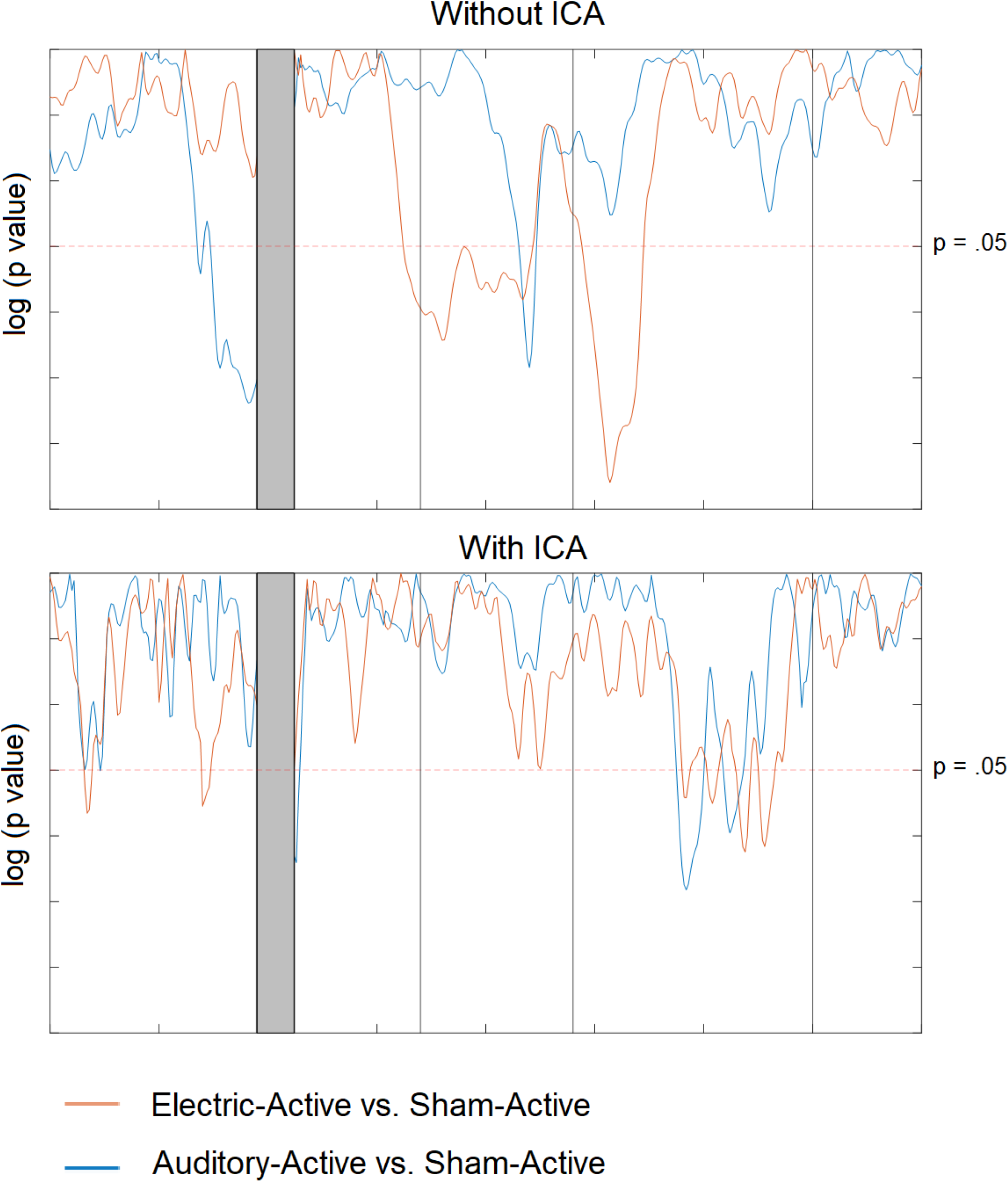
Effects of excluding ICA from the pre-processing pipeline on the comparisons of the correlations between Sham-Active and Electric-Active, and Sham-Active and Auditory-Active.

